# A role of villin-dependent F-actin organization in peroxisome motility in Arabidopsis cells

**DOI:** 10.1101/2025.04.23.650090

**Authors:** Calvin H. Huang, Amanda M. Koenig, Yuh-Ru Julie Lee, Yibo Shi, Jianping Hu, Bo Liu

## Abstract

Actin microfilaments (F-actin) serve as the track for directional movement of organelles in plants. To understand how the F-actin network is employed for the movement of peroxisomes, essential organelles in plant metabolism, we analyzed mutants of three villin (*VLN*) genes, which encode the primary actin-bundling factor and are most actively expressed in vegetative tissues in *Arabidopsis thaliana*. We found that the *vln4* mutation greatly exacerbated the growth and subcellular defects in *vln2 vln3*. Compared to the wild-type cells, the double and triple *vln* mutants exhibit progressive reduction of stable F-actin bundles and rapid remodeling of the fine filaments. The defective F-actin network did not prevent peroxisomes from taking on both rapid and slow movements along the tracks but caused significantly reduced speed of movement and displacement distance of peroxisomes. Using a correlation analysis method, we classified the complex heterogeneous peroxisome movement patterns into clusters reflecting distinct movement directionalities. The *vln* triple mutant had significantly reduced number of peroxisomes with long-range and linear movement. Our results provide insights into how VLN-dependent F-actin organization is coupled with the complex pattern of peroxisome movement.

## INTRODUCTION

The single-membrane organelle peroxisome displays rapid directional movement through the cytoplasm while performing essential functions such as detoxification, β-oxidation, and phytohormone biosynthesis. All these functions likely require peroxisome translocation throughout the cell to scavenge cytotoxins and coordinate with other cellular compartments for versatile metabolic reactions (Sandalio et al., 2021). In mammals, peroxisomes interact with the endoplasmic reticulum (ER), lipid droplets, lysozymes, and mitochondria during metabolic processes like fatty acid catabolism, and defects in peroxisomal distribution and inter-organellar interactions are the underlying cause of several diseases in humans (Wanders et al., 2023). In plants, peroxisomes also exhibit critical interactions with chloroplasts for photorespiration and phytohormone biosynthesis, besides their interaction with the ER and mitochondria (Koenig et al., 2023; Pan et al., 2020). Additionally, peroxisomes participate in functions across the entire expanse of the cell, and their trafficking and spatial arrangement are often linked to the fluctuating demands of the cell during development or under stresses (He et al., 2021). The dynamic distribution of peroxisomes, therefore, requires agile motility along the cytoskeletal filaments and their associated motor proteins (Neuhaus et al., 2016).

In fungi and animals, peroxisome trafficking is primarily governed by microtubules and the associated motors of dynein and kinesins (Covill-Cooke et al., 2021; Salogiannis and Reck-Peterson, 2017). In plants, however, active movement of peroxisomes and other organelles predominately takes place on the F-actin tracks and depends on the motor proteins in the Myosin XI subfamily (Avisar et al., 2008). In *Arabidopsis thaliana*, for example, actin depolymerization or inactivation of multiple Myosin XI genes causes a nearly complete cessation of organelle movement (Mathur et al., 2002; Peremyslov et al., 2010). However, research to date has barely explored the molecular mechanisms that regulate actomyosin-dependent organelle movement in plants (Koenig et al., 2023). In typical differentiated plant cells, F-actin is organized into two distinct populations: long and stable bundles and highly dynamic filaments (Henty-Ridilla et al., 2013). It is unclear whether organelles like peroxisomes employ either or both types of F-actin tracks for their directional movement.

F-actin bundling in plants is primarily mediated by the villin (VLN) family proteins, which were initially isolated from lily pollen as abundant actin-binding proteins (Huang et al., 2015; Yokota et al., 2003). In Arabidopsis, VLN proteins are encoded by five genes and show various degrees of actin bundling capabilities both *in vitro* and *in vivo* (Li et al., 2025). Of the five paralogs, *VLN2*, *VLN3* and *VLN4* are most broadly expressed in vegetative tissues. The *vln2 vln3* double mutants display a reduction in F-actin bundling and altered growth patterns with twisted organs, but no significant compromise in vegetative growth (Bao et al., 2012; van der Honing et al., 2012). Therefore, we hypothesized that VLN4 might function redundantly with VLN2 and VLN3 in mediating organelle movement and the coupled growth processes in plants. Here, we analyzed F-actin organization and peroxisome movement in the mutants of these three Arabidopsis *VLN* genes highly expressed in vegetative tissues. The *vln2 vln3 vln4* triple mutant showed greatly reduced F-actin bundling, increased mesh-like F-actin networks, and gross defects in vegetative growth, which allowed us to interrogate how compromised F-actin bundling impacts patterns of peroxisome movement without disassembling the entire F-actin network. Our analysis of the *vln* triple mutant uncovered significantly reduced movement speed and decreased displacement distance of peroxisomes. We also developed quantitative methods to differentiate features associated with peroxisomal movement modes and correlate such differences with the varied F-actin organization patterns.

## RESULTS

### F-actin organization is deficient in the *vln* mutants

Although peroxisomes are known to move along F-actin tracks by Myosin XI motors in plant cells, it is unknown how the arrangement of the F-actin network facilitates modes of motility, specifically directional movement. We sought to understand the role of actin organization, particularly F-actin bundling, in peroxisome movement. Since the VLN proteins contribute significantly to actin bundling in plants (Huang et al., 2015), we used Arabidopsis *vln* mutants to address how F-actin bundling might be linked to peroxisome movement. Of the five Arabidopsis *VLN* paralogs, *VLN2*, *VLN3* and *VLN4* are most broadly expressed in vegetative tissues according to community-generated transcriptional data (https://bar.utoronto.ca/eplant/). It has recently been reported that VLN2, VLN3, and VLN4 function redundantly in hypocotyl elongation, with VLN2 playing the most prominent role, and that higher-order *vln* mutants display aberrant actin organization in the root and leaf cells (Li et al., 2025). Therefore, we predicted that these three VLN proteins function redundantly to regulate organelle movement and the coupled growth processes.

To determine the role of VLN in peroxisome dynamics, we isolated a *vln4* T-DNA homozygous mutant from the ABRC (Arabidopsis Biological Research Center) and crossed it with the *vln2 vln3* (*vln2/3*) double mutant (Bao et al., 2012). This independently generated *vln2 vln3 vln4* (*vln2/3/4*) triple mutant displayed growth defects (Supplemental Figure 1) similar to what was described recently (Li et al., 2025).

Because of the reported impact of actomyosin-dependent cell growth and the defect of hypocotyl cell elongation in the *vln2/3/4* mutant (Li et al., 2025), we used hypocotyl cells to characterize the actin defects in more detail before investigating peroxisomal dynamics in the mutants. Using live-cell imaging of F-actin labeled by a LifeAct-GFP (green fluorescent protein) fusion protein, we first confirmed that F-actin cable formation is compromised in hypocotyl cells in both the double and triple *vln* mutants (Figure 1A). Compared to the wild-type (WT) cells in which thick, continuous F-actin bundles spanned the cell length, fine F-actin filaments in the *vln* mutant cells were more prominent and organized into mesh-like networks (Figure 1A). In addition, F-actin bundles are less frequently observed in *vln2/3* and completely absent in the *vln2/3/4* triple mutant (Figure 1A). We further quantified the reduction of actin bundles in *vln* hypocotyl cells using the parameter skewness, a standard metric used for assessing F-actin bundling, in which higher fluorescence intensity of the LifeAct-GFP signal in bundled filaments resulted in a higher skewness value (Higaki, 2017; Higaki et al., 2010). The *vln2/3* and *vln2/3/4* mutants exhibited progressively reduced skewness compared to the WT, indicating decreased F-actin bundling in the mutant cells (Figure 1B).

**Figure 1.**
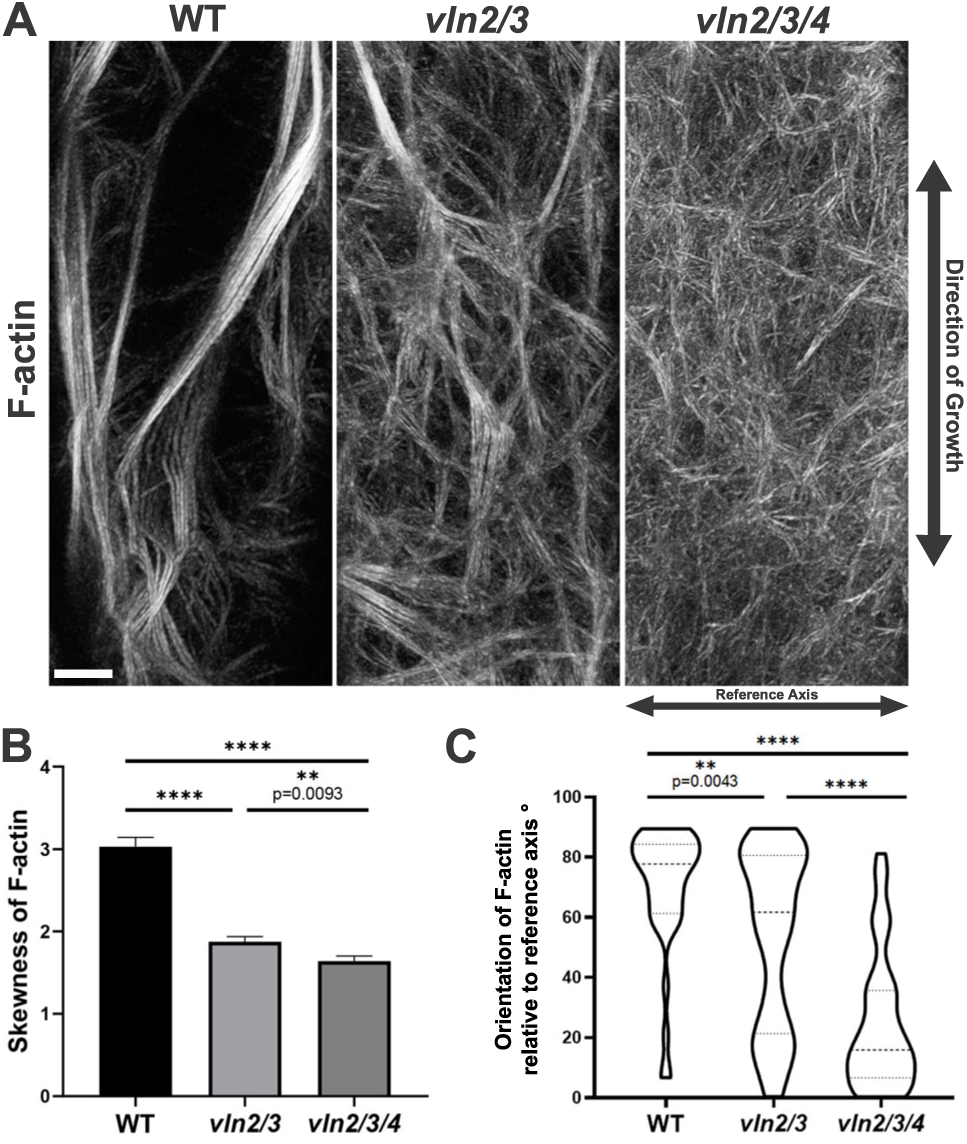
F-actin organization in the Arabidopsis *vln* mutants. (A) LifeAct-GFP-labeled F-actin imaged at the cell cortex of hypocotyl epidermal cells in transgenic wild-type (WT), *vln2/3*, and *vln2/3/4* plants. The WT cell has cortical F-actin in two populations: long, thick bundles and finer filaments in mesh network. The prominent bundles are reduced in *vln2/3* and almost completely lost in *vln2/3/4*. Maximum projections of the LifeAct-GFP images taken over time are shown for all genotypes. (B) The skewness of the F-actin as a measurement of how well the filaments are bundled. Values were obtained from n>35 cells of each genotype. (C) Quantitative assessment of F-actin orientation, determined as the angle relative to the reference axis, where 90° angle is parallel to the direction of growth indicated by double arrows in (A). The mean values are indicated by thick dotted lines. Significance is determined by a two-tailed t-test with p < 0.01 (****). Scale bar: 5 µm.

In WT cells, prominent F-actin cables are often oriented in parallel to the cell elongation axis, likely underlying the recently reported cell elongation phenotype in *vln* mutant hypocotyls (Li et al., 2025). To further assess the deficient F-actin network in the hypocotyl of the *vln* mutants, we quantified the average angle of F-actin filaments, with 0° being in the transverse direction (as a reference axis) and 90° being parallel to the growth direction (Figure 1A). In epidermal cells of the WT hypocotyl, due to the presence of longitudinal bundles along the growth direction, the F-actin filaments were more parallel to the direction of growth, as indicated by a higher average angle (Figure 1C). In the *vln2/3* and *vln2/3/4* mutant cells, however, the overall angles became smaller, concomitant with the decrease in long F-actin cables and the increasingly abundant appearance of mesh networks of F-actin (Figure 1C).

### The dynamic and cell-spanning organization of F-actin is maintained in *vln* mutants

Although *vln* mutants are deficient in actin bundling, they retain a reorganized F-actin network, unlike complete elimination of the filaments by pharmacological means. Therefore, we hypothesized that some actin-dependent organelle motility might be maintained in the mutants. To better understand the specific actin structure available for cargo transport, we further characterized the F-actin network in the *vln* mutants—using stability, coherency, and density as the metrics.

Generally, thick F-actin fibers tend to be stable over time, whereas actin filaments in the mesh network are more dynamic (Henty-Ridilla et al., 2013). Because the F-actin mesh network is pronounced in the *vln2/3* and *vln2/3/4* mutants, we tested whether the altered array organization was associated with rapid actin remodeling in these mutants. Images of F-actin were captured at 5-sec intervals, after which images of different timepoints were pseudo-colored and merged. Noticeably, while F-actin in WT cells predominantly appeared as ‘white’ in the merged images, indicating little, if any, remodeling, the mutant cells showed more dynamic changes in F-actin organization with each timepoint observed in discrete colors (Figure 2A). Further, the white color in the merged images often correlated with thick, long F-actin bundles that are more static than those filaments in the mesh networks in the same WT cells (Figure 2A). In the *vln2/3* and *vln2/3/4* mutant cells, the three distinct colors were more evidently discerned (Figure 2A). Likewise, quantification of F-actin stability showed that a significantly higher proportion of F-actin in the *vln* mutants underwent remodeling than in the WT, with the *vln2/3/4* mutant cells containing significantly fewer F-actin cables than *vln2/3* (Figure 2B). Our data supported previous conclusions that loss of these three VLN proteins severely impaired F-actin bundling and additionally demonstrated the predominance of a diffuse, dynamic mesh network of F-actin in the high-order *vln* mutants.

**Figure 2.**
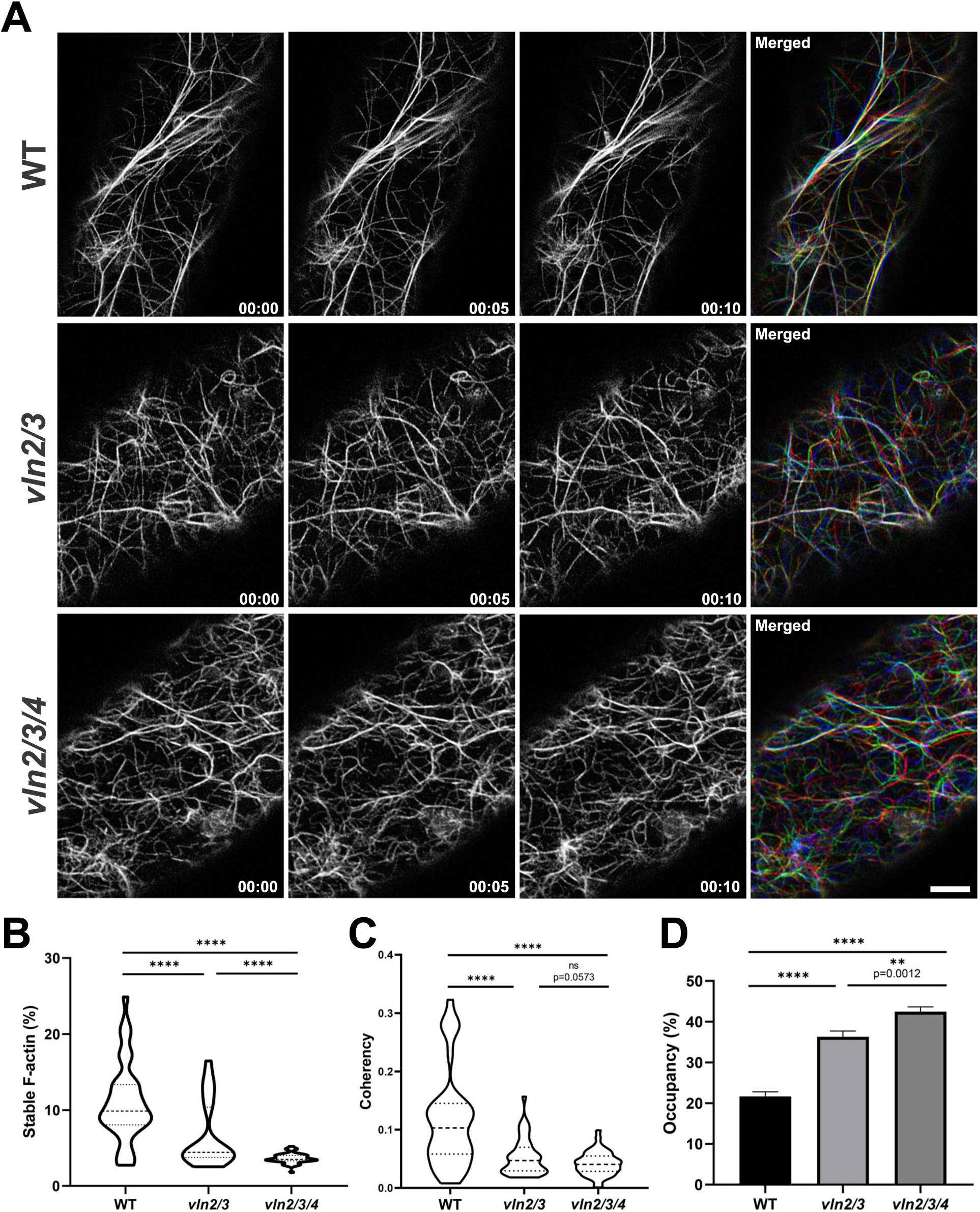
Effects of *vln* mutations on the dynamics of F-actin in Arabidopsis. (A) Three time-lapse images of F-actin taken at 5-sec intervals in the WT, *vln2/3*, and *vln2/3/4* cells. In the merged images, each frame is pseudo-colored (red at 0 sec, green at 5 sec, and blue at 10 sec) to reflect the changes of F-actin filaments where stable filaments with no changes after 10 sec appear in white. WT cells have more stable filaments, whereas the double mutant *vln2/3* has fewer stable filaments and the triple mutant *vln2/3/4* mostly has rapidly remodeling of F-actin filaments. (B) Quantification of the amount of stable F-actin. WT has the highest amount while the *vln2/3* and *vln2/3/4* mutants progressively have less. Values are measured from n>35 cells of each genotype. (C) Coherency of F-actin filaments as a measurement of how F-actin filaments are aligned. Higher coherency values represent more ordered filaments in a preferred direction. (D) Comparison of F-actin density in WT, *vln2/3*, and *vln2/3/4* cells. Density value is determined by the proportion (%) of the cell cortex occupied by F-actin. Significance is determined by a two-tailed t-test with p < 0.01 (****). Scale bar: 5 µm.

Coherency is a measure of the degree of isotropy of the F-actin distribution, where 0 indicates entirely random orientation and 1 indicates uniform orientation, the latter of which is characteristic of cells enriched with actin cables. F-actin filaments in WT cells are organized in a pattern with higher coherence values compared to the presence of more disordered filaments with coherence closer to 0 in the *vln2/3* and *vln2/3/4* mutants (Figure 2C). The decreased coherency in the *vln* mutants suggests the increased occurrence of dispersed and mesh F-actin resulted from the reduced presence of cell-spanning F-actin bundles.

To further characterize the more diffuse, less linear F-actin network in *vln* mutants, we quantified the density of F-actin by measuring the percentage of F-actin occupancy in the cytoplasm. The *vln2/3* cells exhibited significantly higher F-actin occupancy than the WT cells, and the F-actin density in *vln2/3/4* was approximately double that of the WT (Figure 2D). The increased F-actin occupancy suggests that the F-actin network, despite being devoid of obvious bundles, still spans the cortex in the mutants. Therefore, the cytoskeletal apparatus capable of supporting organelle motility across the whole cell seems to persist in the mutants.

To determine if the VLN proteins are associated with F-actin, we expressed a VLN2-GFP fusion protein under the control of its own promoter in the *vln2/3* double mutant. Suppression of the mutant growth phenotype (Supplemental Figure 2A) suggests that VLN2-GFP is functional. When co-expressed with LifeAct-mScarlet-I in tobacco cells, VLN2-GFP was mostly detected on F-actin bundles (Supplemental Figure 2B). Similarly, A VLN4-GFP fusion protein expressed under its native promoter also decorated F-actin bundles marked by LifeAct-mScarlet-I when transiently expressed in tobacco cells (Supplemental Figure 2B).

Taken together, our data showed that in the *vln* mutants, F-actin dynamic polymerization and cell-spanning organization are maintained while only cable formation is deficient. As such, these knockout mutants can serve as an informative genetic tool to study the role of actin organization in peroxisome motility.

### VLN-dependent actin bundling facilitates peroxisome movement

Because of their intact F-actin filaments yet compromised F-actin bundles, the *vln* mutants seemed to be an ideal system to evaluate the specific contribution of actin bundling to peroxisome motility. To determine the impact of actin bundling in peroxisome movement and distribution in Arabidopsis, we transformed WT, *vln2/3*, and *vln2/3/4* plants with constructs for the markers of F-actin (LifeAct-GFP) and peroxisomes (mScarlet-I-SRL). Peroxisomes and F-actin were visualized simultaneously in the hypocotyl cells of plants co-expressing the two fusion proteins. The transgenic *vln* double and triple mutants exhibited progressive deficiencies in F-actin bundling, resembling what was observed earlier in this study (Figure 3A). These defects were concomitant with the progressive aggregation of peroxisomes, suggesting an inability of peroxisomes to be distributed uniformly across the cell in the mutants (Figure 3A). Maximum projections using the time dimensions are shown. It is worth noting that peroxisomes of WT and occasionally in *vln2/3* mutants were able to make long, linear tracks across the cell whereas peroxisomes in *vln2/3/4* moved mostly within a confined area (Figure 3A). Time-lapse images, taken at 5-sec intervals in epidermal cells of the hypocotyl, revealed that peroxisomes maintained motility along the F-actin tracks in both *vln2/3* and *vln2/3/4* mutants (Figure 3A, Movie S1). Moreover, each track of peroxisome movement was heterogenous, with some rapid movements (>1.5 µm/s) as well as more frequently observed slower movements (<0.5 µm/s), similar to what was observed in the WT (Figure 3B), indicating that the mechanisms of actomyosin-dependent peroxisome movement were at least partially maintained in the *vln* mutants.

**Figure 3.**
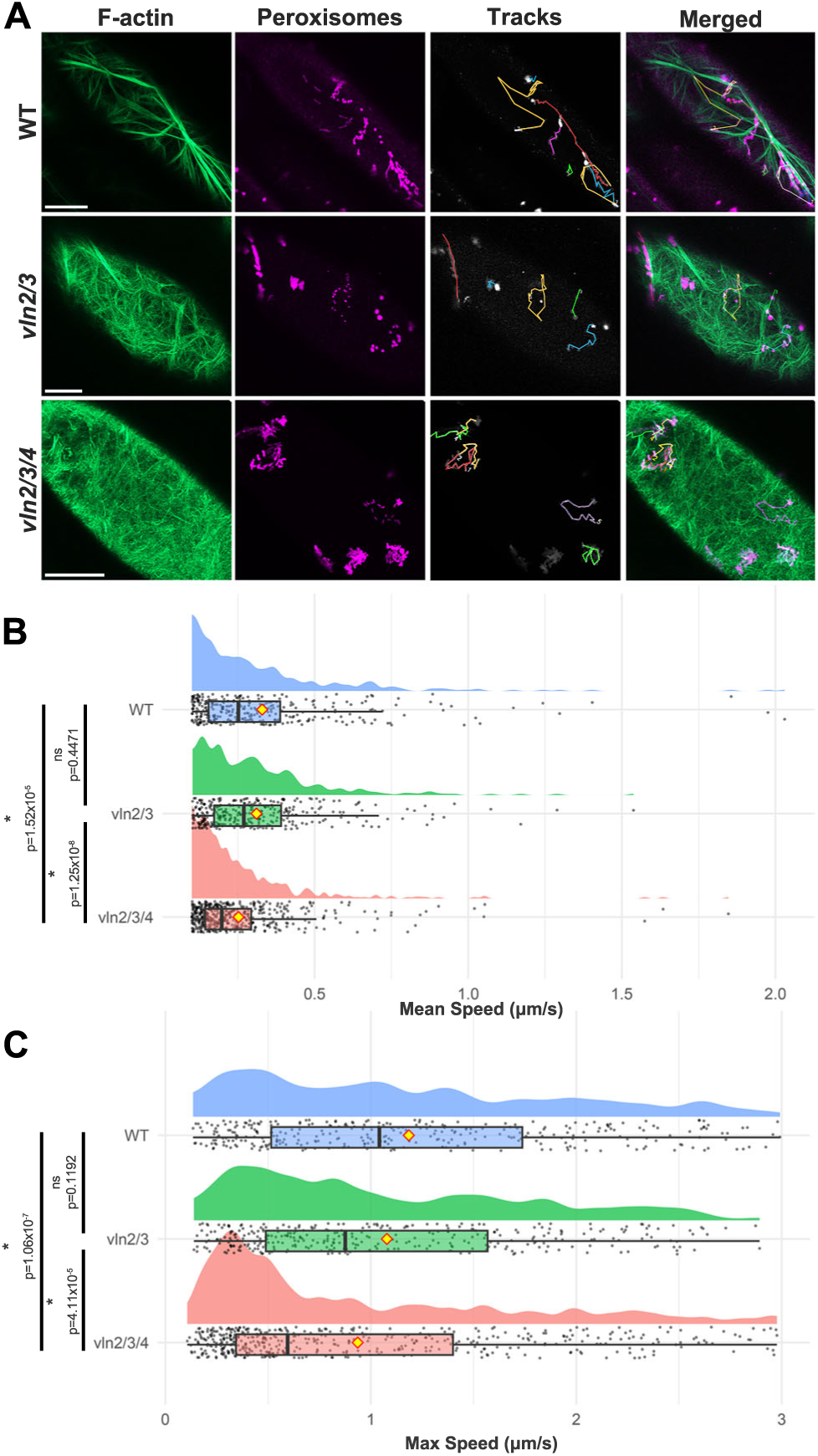
Peroxisome motility is affected in the *vln2/3/4* mutant lacking F-actin bundling. (A) Maximum projections over time of peroxisome and F-actin simultaneously imaged at 5-sec intervals. In the *vln2/3* and *vln2/3/4* mutant cells that have reduced long F-actin bundles, peroxisomes show aggregation, impaired distribution, and reduced linear movement. (B-C) Quantitative assessments of average peroxisome speed and maximum speed in WT and *vln* mutants. Values are measured from n>300 peroxisome tracks from each genotype and plotted along a raincloud plot with a histogram on top and a boxplot below. Yellow diamonds represent the mean position. Statistical significance is determined by a Wilcoxon rank sum test with p-values shown in the figures.

To examine the role of VLNs in peroxisome motility in more depth, we compared the speed of peroxisome movement among WT, *vln2/3* and *vln2/3/4*. Despite variations in instantaneous speed during imaging duration, calculation of the mean speed of each peroxisome movement revealed that only the severe inhibition of F-actin bundling in *vln2/3/*4 caused a significant decrease in the mean speed, while intermediate bundling in *vln2/3* was sufficient to maintain peroxisome speed comparable to that of the WT (Figure 3B). These data suggest that peroxisome movement is facilitated by both discrete F-actin filaments and densely bundled actin cables.

Next, we wanted to exclude peroxisomes that were potentially unable to associate with F-actin via myosin from our calculation and only assess the movement of peroxisomes already associated with myosin motors. To achieve this goal, we calculated the maximum speed of peroxisome movement, defined as the highest instantaneous speed recorded between two consecutive timepoints. Again, the maximum speed distributions were not significantly different between WT and *vln2/3*, but were more frequently observed at lower values in *vln2/3/4* (Figure 3C). We therefore conclude that actin dynamics and organization influence peroxisome velocity and distribution.

### Fast, directional peroxisome movement and distribution are facilitated by F-actin cables

Since the heterogenous pattern of peroxisome motility shifted towards slower speeds in the absence of longitudinal actin bundles, as shown in the *vln2/3/4* triple mutants, we hypothesized that F-actin cables specifically facilitate fast, linear movement, enabling uniform peroxisome distribution across the cell. To test this, we evaluated total moving distance, displacement, straight-line speed, overall linearity of movement, and directional change rate of peroxisomes when they deviated from the prior trajectories (Figure 4A-E).

**Figure 4.**
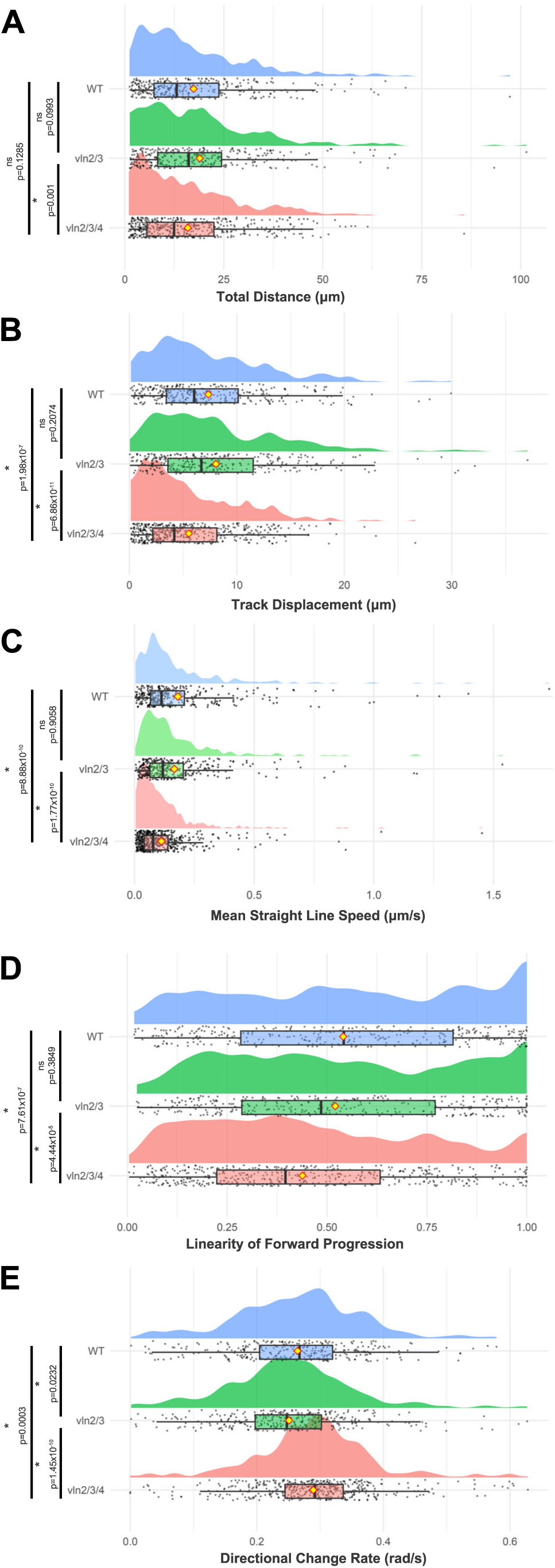
Quantitative comparison of peroxisome tracks in the WT, *vln2/3* and *vln2/3/4* mutant cells. Parameters include total distance (A), track displacement (B), average straight-line speed (C), linearity as a ratio of straight-line speed to track mean speed (D), and directional change rate in radians (rad)/s (E). Values are obtained from n>300 peroxisome tracks from each genotype and plotted in a raincloud plot with a histogram on top and a boxplot below. Yellow diamonds represent the mean values. Significance is determined by a Wilcoxon rank sum test with p-values shown in the figures.

Measurement of the total moving distance revealed that most peroxisomes traverse similar distances with no significant difference between mutant and WT cells (Figure 4A). However, peroxisomes in *vln2/3/4* cells have a significantly shorter displacement, measured as the shortest distance between the start and end points of a tracked peroxisome (Figure 4B). Consistently, the triple mutant also exhibited reduced displacement distance, whose value was normalized by track duration and represented by “straight-line speed” that accounts for any variations of movements tracked per peroxisome (Figure 4C). Together these data suggest that, while peroxisomes still move in short bursts that sum to a similar distance observed between *vln* mutants and WT, these organelles do not successively travel as far distance in the *vln2/3/4* mutant, which lacks prominent F-actin bundles or cables, when compared to WT or *vln2/3*. In fact, peroxisomes in *vln2/3/4* mutant cells changed direction more frequently than those in the WT cells. Motility in *vln2/3/4* was overall less linear with a lower ratio of straight-line speed to track mean speed (Figure 4D). However, peroxisomes in the triple mutant more frequently change directions (Figure 4E), likely because their movement predominantly relied on the dynamic F-actin mesh network rather than stable, longitudinal bundles as tracks.

To better assess the heterogenous motility pattern of peroxisomes, we employed a Pearson’s correlation matrix to conduct pairwise analysis of the influence of the aforementioned motility parameters on each other within individual genotypes. The *vln* mutants showed correlations comparable to WT between average velocity, maximum speed, total distance and track displacement. By contrast, correlations between parameters that account for directional movement (straightforward speed, linearity, and directional change rate) varied across genotypes (Supplemental Figure S3A). We then plotted the data of straight-line speed, linearity, and directional change rate from all genotypes in a 3-dimensional scatterplot and used k-means clustering to group organelle motility according to these multidimensional criteria (Figure 5A). Using the Elbow Plot method (Supplemental Figure S3B), the data were grouped into 3 clusters: high straightforward speed and linearity but low directional change rate as Cluster 1; intermediate straightforward speed, linearity, and directional change as Cluster 2; and low straightforward speed and linearity but high directional change rate as Cluster 3. Cluster 1 is enriched for peroxisomes that rapidly and directionally traverse the cell with minimal turns, while Cluster 3 represented peroxisomes that move in short bursts with frequent changes in direction. Interestingly, WT peroxisomes exhibited uniformly heterogeneous movement across the three clusters, with roughly one third of the tracked movements represented in each cluster (Figure 5A). However, in the *vln2/3* mutants, a smaller proportion (29.6% in Cluster 1) of peroxisomes move directionally compared to 32.8% in the WT, and the peroxisome motility profiles shifted towards intermediate (Cluster 2) and short burst (Cluster 3) movements (Figure 5A). In the *vln2/3/4* triple mutant, the reduction in fast, linear peroxisome movements was further exacerbated (20.6% in Cluster 1), with more peroxisomes moving in short bursts (39.8%, Cluster 3) compared to 30.5% in WT (Figure 5A). This shift in cluster populations across genotypes suggests that a lesser proportion of peroxisomes move quickly and linearly when F-actin bundling is deficient in the *vln* mutants.

**Figure 5.**
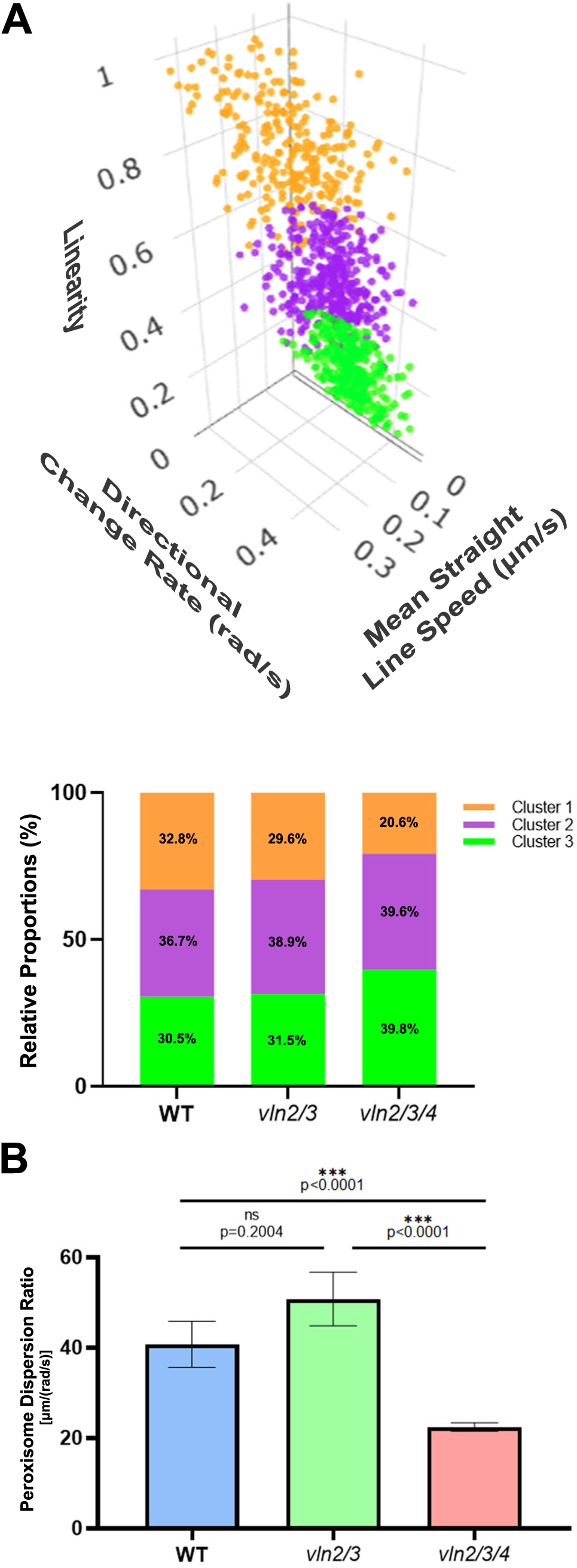
Multivariate clustering and dispersion analysis distinguish peroxisome movement behaviors in WT and *vln* mutants. (A) Pairwise correlation analysis of peroxisome movement in directional change rate, average straightforward speed, and linearity. Formation of three clusters resulted from the pairwise correlation analysis. Proportion of peroxisome movement patterns is represented by the three clusters in WT, *vln2/3*, and *vln2/3/4* cells. (B) Peroxisome dispersion ratios used to quantify peroxisome distribution, as reflected by the rate of track displacement and directional change [µm/(rad/sec)]. Peroxisomes in WT and *vln2/3* are able to travel long distances before changing directions, shown by higher ratio values. Conversely, peroxisomes in *vln2/3/4* show more confined movements within a localized region, evidenced by lower ratio values when compared to WT and *vln2/3*. Significance is determined by a two-tailed t-test with p-values shown in the figures. Values are measured from n>300 peroxisomes for each genotype.

To assess whether peroxisome distribution was altered in *vln* mutants, we determined the ratio of track displacement to directional change rate. A higher ratio indicates greater dispersion, as it reflects directed movement for a greater distance (*i*.*e*., more displacement with few directional changes). In contrast, a lower ratio reflects movement within a more confined region, representing frequent directional changes with little displacement. In WT and to some extent *vln2/3* mutants, long and stable F-actin cables serve as the track for peroxisomes to be dispersed throughout the cell. However, the absence of such cables in *vln2/3/4* would limit peroxisomes to move predominantly within localized regions. In agreement with this prediction, the outcome of our quantification assessments indicated that the WT and *vln2/3* cells exhibited higher average dispersion ratios [40.76 ± 5.10 and 50.8 ± 5.92 µm/(rad/sec), respectively] than those of *vln2/3/4* [22.48 ± 0.97 µm/(rad/sec)] (Figure 5B). These findings suggest that peroxisome movement becomes more confined within the actin mesh network in the triple mutant.

Taken together, our results showed that densely bundled F-actin cables are necessary for rapid, directional movement of peroxisomes, which likely enable the proper subcellular distribution of peroxisomes required for their diverse metabolic functions.

## DISCUSSION

In this work, we generated *vln2/3/4* triple mutant to investigate the collective roles of three F-actin bundling VLN proteins in the dynamics and organization of the F-actin network. Our quantitative analysis of peroxisome motility in the WT and *vln* mutants showed that VLN-dependent F-actin organization is required for rapid directional movement of peroxisomes in the epidermal cells of Arabidopsis hypocotyls.

Plant VLN proteins are grouped into three classes, in which VLN4 belongs to class III while VLN2 and VLN3 are in class II (Khurana et al., 2010). Consistent with what was reported recently (Li et al., 2025), we found loss of VLN4 to greatly enhance defects in the *vln2/3* double mutant. Our results collectively indicate that the functions of these three VLN proteins, despite being in two different classes, converge towards stabilizing F-actin and promoting high-order F-actin bundle formation in vegetative tissues. However, when the three VLN proteins were examined individually without an F-actin marker, they exhibited different extents of localization on F-actin-like structures in different tissues (Li et al., 2025). Here we showed that both VLN2 and VLN4 proteins had pronounced association with prominent F-actin bundles but less so with fine filaments (Supplemental Figure S2B). Although VLN proteins exhibit common actin-bundling activities in vitro, it will be interesting to determine by live-cell imaging whether these proteins co-localize and if so whether their localization may be augmented by each other *in vivo*. More importantly, we wish to learn how they perform cooperative functions in bundling and/or stabilization of F-actin filaments *in vivo*. Surprisingly, additional mutations in two other *VLN* genes encoding class I and III VLN proteins respectively did not exacerbate the growth phenotypes in the *vln2/3/4* triple mutant (Li et al., 2025), indicating that the three VLN proteins examined in our study play the primary role in F-actin bundling.

Previous phenotypic analysis of myosin mutants clearly demonstrated that directional movement of organelles like peroxisomes is dependent on Myosin XI motors (Peremyslov et al., 2010; Perico and Sparkes, 2018). It is intriguing how specific organelles can be actively transported at such a wide range of speeds. We hypothesize that the heterogeneity of peroxisome motility can be brought about by different Myosin XI motors that exhibit different motile activities (Haraguchi et al., 2016). When ectopically expressed in a heterologous system, myosin truncations including the DIL (dilute) domain from at least 7 of the 13 Arabidopsis Myosin XI motors exhibit peroxisome-binding activities (Sattarzadeh et al., 2011). However, it is yet to be determined whether the full-length myosin proteins are associated with the organelle when expressed under their native promoters. It is also possible that multiple motors can bind to the same organelle to increase the movement processivity so that it can be transported for long distance, especially in giant cells of organisms like *Chara corallina* that demonstrate long-distance transport of organelles at great velocities (Shimmen and Yokota, 2004). Varied number of active motors on each peroxisome, which is still hard to quantify *in vivo*, likely contributes to the heterogeneity in transport velocities as well.

While Myosin XI motors function as the drive of organelle motility with significant influence on the speed and travel distance, we showed that F-actin organization was critical for the diversity of organelle movement. When VLN-dependent actin bundling is deficient, fewer peroxisomes reached maximum speeds (Figure 3B), and the linearity of peroxisome movement was impaired (Figure 4). Golgi and mitochondria show similar shifts in motility according to the natural variation in F-actin architecture across the root elongation zone (Akkerman et al., 2011). The variation in F-actin networks throughout cell development may support the proper distribution of organelles. For example, longitudinally oriented, prominent F-actin cables are found in plant cells during interphase and are often considered the primary tracks of long-distance transport of organelles for their robust anisotropic expansion (Garcia-González and van Gelderen, 2021). Our analysis of the *vln2/3/4* triple mutant revealed that fine filaments of F-actin are capable of facilitating directional transport of peroxisomes but insufficient to maintain the robust directional movement that takes place on VLN-dependent actin cables. However, the long, stable F-actin bundles exhibit great advantages of promoting rapid, directional movement of the organelles. Compromises of such rapid, directional movement have a profound consequence in cell expansion that presumably is dependent on rapid deployment of organelles across the cytoplasm.

In addition to developmentally related roles, both organelle distribution and VLN-dependent actin bundling have been linked to cellular responses to biotic stresses like bacterial infection. For example, inoculation of virulent bacteria induces bundling of F-actin in Arabidopsis leaf cells (Lu et al., 2020; Shimono et al., 2016). It will be very interesting to test whether changes in F-actin organization and/or organelle transport brought about by *vln* mutations influences their susceptibility to pathogenic microbes.

We observed highly heterogeneous peroxisome movement, the significance of the which remains unclear. Because peroxisomes often exchange metabolites with other organelles for their function, for example with mitochondria and chloroplasts during photorespiration, it is likely that peroxisome motility is coupled with the position of other organelles to facilitate inter-organelle communication. In fact, chloroplast-mitochondrion-peroxisome interactions increase in response to high light (Jaipargas et al., 2016), which induces higher rates of photorespiration, and mitochondria exhibit different motility patterns before and after being associated with chloroplasts (Oikawa et al., 2021). It would be interesting to determine whether certain movement patterns are unique for peroxisomes physically interacting with other organelles.

Additionally, peroxisome velocities may vary across the cytoplasm and are coupled with localized spaces with different physiological conditions. Lastly, various intrinsic and extrinsic factors may cause distinct velocities and directionalities of peroxisomes, which may be enabled by VLN-dependent F-actin bundling for efficient motility and distribution.

## METHODS AND MATERIALS

### Plant genotype and growth

*Arabidopsis thaliana* plants were grown in soil in a growth chamber at 21°C with cycles of 16hr light and 8hr dark. *Arabidopsis* seedlings that were used for imaging were sterilized and sowed onto ½ Murashige & Skoog basal media with 0.8% phytagel. Seeds in solid media were cold stratified for 2 days then moved into the growth chamber for 6 days before imaging. All *Arabidopsis thaliana* mutants and WT plants are in the Col-0 genetic background. Floral dipping method was used to deliver transgenes into the plant, following established protocols (Clough and Bent, 1998).

Mutants of *VLN4* were isolated from seed stock of SALK_109099C (Arabidopsis Biological Resource Center, Ohio State University), which carried a T-DNA insertion in the 12^th^ intron of the *VLN4* gene, by using the primers 109099RP (5’-CAT TCA AAA ACT CTA TGC TGT TGG-3’) and LBb1.3 (5’-GGA TTT TGC CGA TTT CGG AAC CAC-3’) to detect the T-DNA insertion and 109099LP (5’-TAT ATC TCT GTC ACC TGC CAA AAG-3’) and 109099RP to detect the wild-type allele. The *vln2/3* double mutant, generally provided by Prof. Shanjin Huang, was described in an earlier publication (Bao et al., 2012).

### Plasmid construction

To produce an expression vector carrying cassettes for expression of both mScarlet-I-PTS1 and LifeAct-GFP, we first had the codon-optimized mScarlet-I-encoding DNA fragment synthesized commercially (Integrated DNA Technologies, Inc.). The SRL-coding sequence was added to the 3’ end by PCR using primers 5’-GTA CCG AAT GGT GAG CAA GGG AGA AG-3’ and 5’-CGG CGC GCC TTA TAG CCG GCT TTT ATA CAG TTC GTC CAT ACC GCC G-3’. The resulting fragment was inserted into a plasmid containing the promoter sequence of EF1α (AT1G07920). Separately, the LifeAct-coding sequence was synthesized and inserted into pENTR4 (Thermo Fisher), to generate a plasmid expressing LifeAct-GFP under the viral 35S promoter by Gateway LR cloning with the pGWB5 plasmid (Nakagawa et al., 2007). The resulting plasmid was linearized by *Apa*I, after which the DNA fragment for EF1α(p)::mScarlet-I-PTS1 (SRL), amplified by the primers 5’-GCG ACA ATA TGA TCG GGC CTC CGG TGT CAA TCT CAG G-3’ and 5’-TGT GGA CGC CGG GCC TTA TAG CCG GCT TTT ATA CAG TTC G-3’ from the plasmid described above, was inserted by Gibson reaction to generate the plasmid pLB187 used for transformation.

Plasmids were transformed into the Agrobacterium strain GV3101, prior to floral dipping in the WT, *vln2/3* and *vln2/3/4* plants.

### Confocal microscopy

After germination and growth in the chambers, Arabidopsis seedlings were mounted in water for imaging. Live-cell imaging was performed on an LSM980 laser scanning confocal microscope with the Airyscan2 function using a 40X water immersion objective and standard excitation and emission settings for EGFP and mCherry by the manufacturer (Carl Zeiss). Single frame images of F-actin were used to measure orientation, coherency, density and skewness. For measurements of F-actin stability, time-lapse images with 5sec/frame were obtained. Similarly, to image peroxisome movement, both markers for F-actin and peroxisomes were simultaneously imaged with a time-lapse of 5sec/frame. At least 3 independent transformant lines were imaged per genotype. In Figure 1A and 3A, time-lapse movies were presented in maximum intensity projection of images collected over time.

### Quantification of F-actin

Images were first rotated such that the direction of hypocotyl growth is along the vertical axis, then thresholded, binarized and skeletonized for F-actin quantification (Higaki, 2017). A region of interest was drawn and selected for quantitative analysis. The ImageJ plugin OrientationJ was used to measure orientation and coherency (Püspöki et al., 2016). Skewness and density were measured using the LPX plugin package (Higaki et al., 2010; Schneider et al., 2012).

In measuring the percentage of stable F-actin, F-actin was imaged for 10 sec at 5 sec/frame, after which the first (time zero) and the last (10 sec) frames were merged. Under the reasoning that stable F-actin filaments should still be present or unchanged in position after 10 sec, a stable filament from the last frame should colocalize with itself from the first frame. The merged image was thresholded for colocalized filaments. The percentage of stable F-actin was calculated using the ratio of colocalized F-actin signal to the total F-actin signal.

### Quantification of peroxisome motility and distribution

Quantitative measurement of F-actin-dependent peroxisome motility was carried out by imaging F-actin and peroxisomes simultaneously with 5-sec intervals. Peroxisomes not associated with F-actin were not measured. Peroxisomes were tracked using the ImageJ plugin, TrackMate (Ershov et al., 2022). Given that the speed of slow moving peroxisomes is usually ≥0.2 µm/sec (Mathur et al., 2002), we excluded peroxisomes moving at velocities below 0.1 µm/sec as those appear to be more likely due to Brownian movements. We also manually reevaluated the tracks with velocities <0.1 µm/sec to confirm that these movements appeared oscillatory. The output track data summarized the overall movement of individual peroxisomes during the imaging duration, and provided the following track features: (1) mean speed for the average speed values during every 5-sec intervals along a track; (2) maximum speed for the highest instantaneous speed between two consecutive timepoints along a track; (3) straight-line speed, calculated as track displacement divided by track duration; (4) total distance traveled; (5) track displacement; (6) linearity of forward progression for the ratio between straight-line speed and track mean speed, where completely linear tracks would have a value of 1; and (7) directional change rate calculated as the average angle between consecutive timepoints along a track divided by the track duration.

To quantify peroxisome distribution, a dispersion ratio was calculated by having the track displacement (µm) divided by the directional change rate (rad/s). A higher ratio represents peroxisomes that travel long distances with few turns, which would result in peroxisome dispersion across the cell. A lower ratio represents a peroxisome that turns frequently and moves within a small region.

## Acknowledgements

We thank Dr. Shanjin Huang for providing the *vln2/3* mutant seeds and Dr. Tsuyoshi Nakagawa for sharing the pGWB plasmids, Dr. Felicia L. Peng and Dr. Chenxin Li for their advice and comments on this work. This study was made possible in part through access to the MCB Light Microscopy Core Facility with training and support by Dr. Thomas Wilkop.

## Competing interests

The authors declare no other competing or financial interests.

## Author contributions

Conceptualization: C.H.H., Y.-R.J.L., J.H., B.L.; Methodology: C.H.H., A.M.K., Y.-R.J.L., J.H., B.L.; Investigation: C.H.H., Y.-R.J.L.; Formal analysis: C.H.H., A.M.K., Y.-R.J.L.; Visualization: C.H.H., Y.-R.J.L.; Data curation: C.H.H.; Supervision: B.L., J.H.; Project administration: B.L.; Funding acquisition: J.H., B.L.; Writing (original draft): C.H.H., A.M.K.; Writing (review and editing): C.H.H., A.M.K., Y.-R.J.L., J.H., B.L.

## Funding

This work was supported by grants from the U.S. National Science Foundation (2148207 to B.L & 2148206 to J.H). B.L. was also sponsored by the U.S. Department of Agriculture (USDA)-the National Institute of Food and Agriculture (NIFA) under an Agricultural Experiment Station (AES) hatch project (CA-D-PLB-2536-H). A.M.K was partially funded by a grant from the Chemical Sciences, Geoscience and Biosciences Division, Office of Basic Energy Sciences, Office of Science, U.S. Department of Energy (DE-FG02-91ER20021) to J.H. The Zeiss LSM 980 microscope used in this study was purchased with a grant (S10OD026702) from the U.S. National Institutes of Health.

## Data and resource availability

All data acquired from this study are included within the main text and Supplementary information.

## Notes

### Competing Interest Statement

The authors have declared no competing interest.

### Summary of Updates

Figure 2 revised; Figure 5 revised

