## Supplemental Figure 1-3 for "A role of villin-dependent F-actin organization in peroxisome motility in Arabidopsis cells"

**Huang et al., 2025. Supplementary Figures**

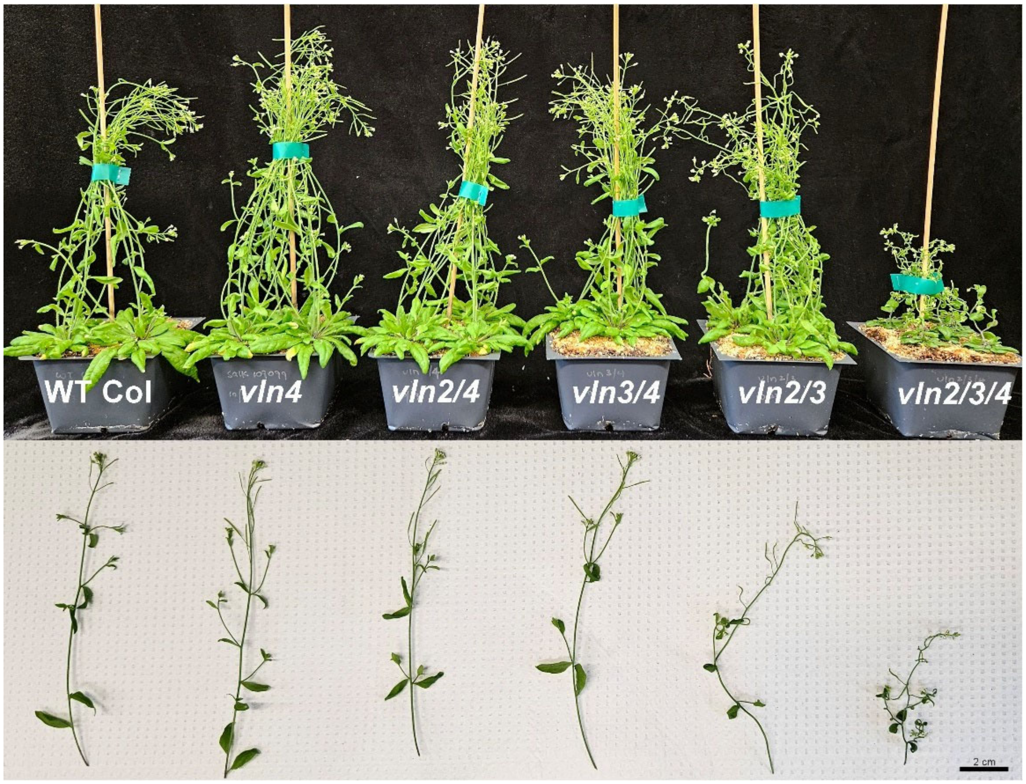


**Supplementary Figure 1.** The *vln4* mutation enhances phenotypes of the *vln2/3* double mutants. Phenotypes of the adult plants (top) and inflorescence branches (bottom) of the wild-type (WT, Col-0), and single, double, and triple mutants of *vln4, vln2/4, vln3/4, vln2/3, and vln2/3/4*, are compared. *The vln4, vln2/4, vln3/4* mutants grow indistinguishably from the WT. The *vln2/3* double mutant shows altered growth postures with undulating organs as represented by twisted inflorescent stems but retains its robust vegetative and reproductive growth. The *vln2/3/4* triple mutant has greatly retarded growth phenotypes including dwarfism with exacerbated posture defects of twisted organs. Scale bar, 2 cm.

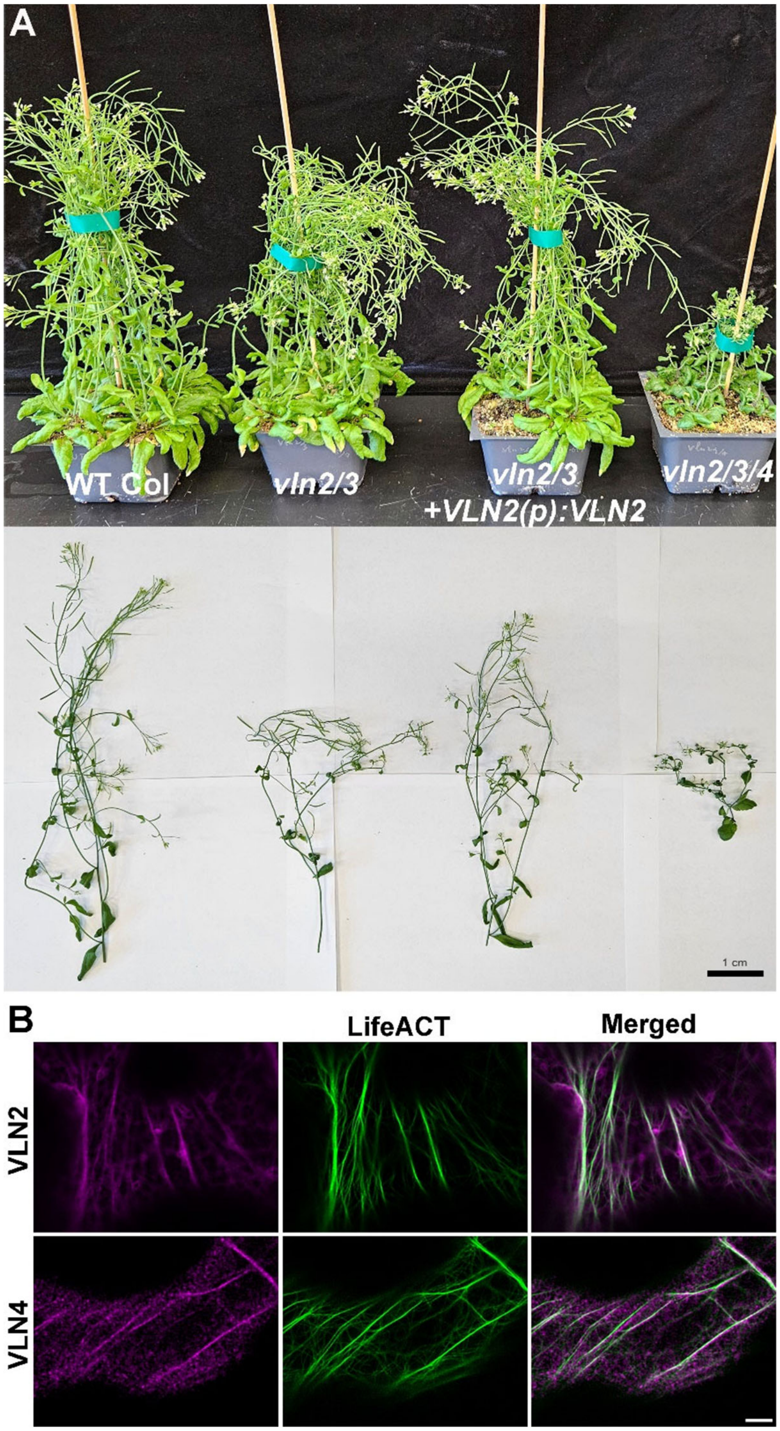


**Supplementary Figure 2.** VLN proteins decorate F-actin bundles. (A) Expression of VLN2-GFP suppresses the *vln2/3* phenotype of postures seen in adult plants (top) and inflorescence branches (bottom). (B) In *N. benthamiana* leaf cells, both VLN2-GFP and mCherry-VLN4 fusion proteins colocalize with F-actin bundles revealed by the LifeAct-mScarlet-I and LifeAct- GFP fusion proteins, respectively. Scale bars, 1 cm (A) and 5 μm (B).

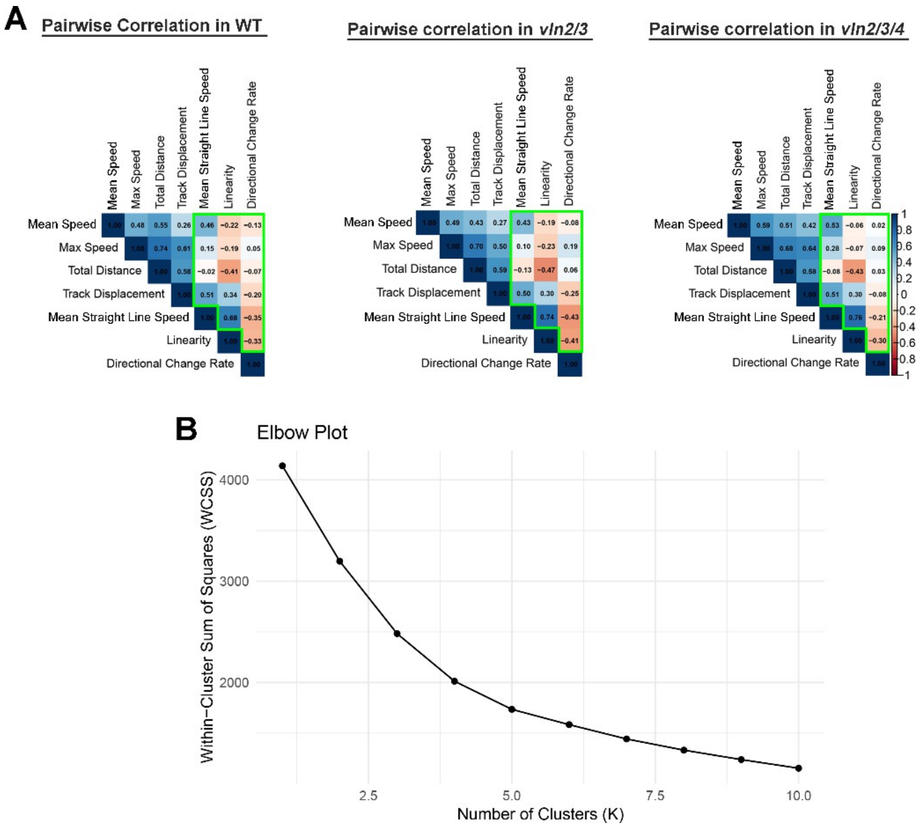


**Supplementary Figure 3.** Analysis of peroxisome movement by pairwise correlation analysis. (A) Pearson's pairwise correlations between the seven measured parameters of peroxisome tracks are determined in the wild-type (WT), the *vln2/3* double mutant, and the *vln2/3/4* triple mutant. Weighted blue colors represent positive correlations and red colors represent negative correlations. (B) An elbow plot is used to determine the appropriate number of clusters to describe the peroxisome tracks. Inflection point is around 3 so that the datapoints are clustered into 3 groups.
